# Multiple Dimerizing Motifs Modulate the Dimerization of the Syndecan Transmembrane Domains

**DOI:** 10.1101/2019.12.31.891929

**Authors:** J. Chen, F. Wang, C. He, S-Z. Luo

## Abstract

Syndecans(SDCs) are a family of four members of integral membrane proteins, which play important roles in cell-cell interactions. Dimerization/oligomerization generated by transmembrane domains (TMDs) appear to crucially regulate several functional behaviors of all syndecan members. The distinct hierarchy of protein-protein interactions mediated by the syndecan TMDs may give rise to considerable complexity in the functions of syndecans. The molecular mechanism of the different dimerization tendencies in each type of SDCs remains unclear. Here, the self-assembly process of syndecan TMD homodimers and heterodimers was studied in molecular details by molecular dynamics simulations. Our computational results showed that the SDC2 forms the most stable homodimer while the SDC1 TMD dimerizes weakly, which is consistent with previous experimental results. Detailed analysis suggests that instead of the conserved dimerizing motif G8XXXG12 in all four SDCs involved in homo- and hetero-dimerization of SDCs, the G3XXXA7 motif in SDC1 competes with the interface of G8XXXG12 and thus disturbs the SDC1 involved dimerization. The SDC3 which contains a G9XXXA13 motif, however, forms a more stable dimer than SDC1, indicating the complexity of the competing effect of the GXXXA motif. As GXXXG and GXXXA are two common sequence motifs in the dimerization of helices, our results shed light on the competing effect of multiple dimerizing motifs on the dimerization of transmembrane domains.

## Introduction

Extracellular ligand recognition plays a vital role in cell signaling and cell-cell interactions. Most cell surface receptors are comprised of the ectodomain, transmembrane domain (TMD) and a cytoplasmic domain[1]. While a lot of studies focus on the ectodomains, which are responsible for the interactions with extracellular ligands, and the cytoplasmic domains, which stimulate intracellular functions[2], less information is provided about the molecular mechanism of the role of TMDs in signaling pathways. Syndecans (SDCs) are a family of four members of integral membrane proteins that functions as binding to various ligands in extracellular matrix through covalently bonded heparan sulfate, acting as co-receptors of heparan sulfate[3]. Syndecan are involved in cellular proliferations, cell-cell adhesion, discrimination, migrations and also wound healing by growth factor production[4]. The four members of syndecan are named as Syndecan-1 (SDC1), Syndecan-2 (SDC2), Syndecan-3 (SDC3) and Syndecan-4 (SDC4)[5]. All members have variable lengths of amino acids, while in humans SDC3 is the longest member.

In present, apart from the cytoplasmic domain of syndecan-4, the 3D structure of syndecans are poorly characterized, which hinders our understanding of the molecular mechanisms underlying syndecan functions that are mediated by numerous interactions.[6] TMDs of all members of the syndecan family is now considered as the pivotal objective of current curative drugs[7]. TMDs play a signaling role between the linking of extracellular and cytoplasmic domains by initiating cytosolic activities via receptor clustering and anatomical change also occurs in its structural compositions when the ectodomain interacts with outer matrix[8]. There have been quite a few experimental studies on the structures and also the roles of different syndecan members in cell signaling, functioning, regeneration and cellular interactions[9–11]. All of the syndecan members are adequate for SDS resistant dimerization even in the absence of both ectodomain and cytoplasmic domain, indicating that TMD is the essential component for the dimerization.

However, the molecular mechanism of the dimerization/oligomerization of different syndecan members still requires further illustration. The aim of our work was to broaden current knowledge of dimerization/oligomerization of the TMDs of all syndecan members. Here, we present Molecular Dynamic (MD) simulation studies on the dimerization/oligomerization of the TMDs of all syndecan members, which agrees with recent reported experimental studies [12] that SDC2 TMD has the strongest tendency of dimerization comparing to others.

## Statement of Significance

Through CGMD it was revealed that in addition to the GXXXG motif present in the transmembrane region of syndecans, the GXXXA motif is also involved in mediating the dimerization between syndecans. The location of the GXXXA motif in the helix chain significantly affects the stability of the dimers.

## Methods

### Unrestrained CG Simulations

All the simulations were performed on the GROMACS-5.1.2[13]. The initial CG structure of transmembrane proteins was firstly generated by Pymol[14–16]. The initial distance of the two transmembrane proteins was set to 5nm. Combined with the treatment of the periodic boundary and the idea of saving the box space, we placed the protein on the diagonal of a box with a size of 7.1×7.1×6.9nm^3^. Charmm-gui[17–19] (http://www.charmm-gui.org/?doc=input/mbilayer) was used to generate files for MD. The martini22p[20, 21] force field and the lipid type DPPC[22–24] bilayer membranes were used. The terminal of protein, which we were dealing with, was neutral. The ion NaCl concentration was 150mM.

After an initial 6000 steps of steepest descent energy minimization, the NVT equilibration was conducted using the v-rescale coupling methods in 200 ns, with a reference temperature of 303.15 K and a time constant of 1.0 ps. The NPT equilibration was conducted by the Parrinello-Rahman[25] coupling methods, by using a coupling constant of 12 ps, a compressibility value of 4.5 ×10^−5^ bar^−1^, and a reference pressure of 1.0 bar. During the equilibration, the isotropic pressure coupling type was applied. The backbones were restricted by a harmony restraint force of 1000 kJ·mol^−1^·nm^−2^ in three dimensions. The electrostatic interactions utilized a dielectric constant of 15. The Lennard-Jones and electrostatic interactions were shifted to zero between 0.9 to 1.2 nm and 0 to 1.2 nm, respectively. After the equilibration accomplished, the position restraint was removed and the initial velocity was generated. The integration time step was set as 20 fs and the pair list was updated every 10 steps. The simulation production ran for a duration of 3 μs. All the presentations were produced by VMD[26] software.

### PMF Calculations

The PMF was calculated using the umbrella sampling[27] technique. Twenty independent simulations, each of 500 ns in duration, were performed for PMF calculation, i.e., the simulations corresponding to the 20 windows were run such that the initial configuration of one simulation did not depend on the outcome of the preceding simulation. Each independent simulation corresponds to a different separation distance between the center of mass of each peptide. To generate the individual window configuration, a position restraint of 1000 kJ·mol^−1^·nm^−2^ was set on the fixed peptide and a pulling force of 800 kJ·mol^−1^·nm^−2^ was added on the pulling peptide at the side of the fibril. A pulling rate of 0.01 nm/ps was used and the pulling time was 500 ps. A distance gap of 0.2 nm was used to select the output configurations to ensure the well-overlapping of the configuration samplings in the following step. After the separated configurations were obtained, all of them went through an NPT equilibration, maintaining the pulling force but set the pulling rate as 0. Then a simulation in 1.0 μs was conducted for each window. The PMF profile was calculated using the weighted histogram analysis method (WHAM) [28].

### The free binding energy Calculations

G_mmpbsa was used to calculate the binding energy of the TMDs of syndecans. The MM, PB, and SA[29] energy values or all energies according to their objective was obtained. G_mmpbsa also gives residue wise contribution to the total binding energy.

## Results and Discussions

### The homodimerization of SDC TMDs

To model the dimerization of syndecan TMDs, two CG helices of syndecans TMDs were inserted in a parallel orientation relative to one another in a preformed DPPC bilayer (Fig S1 left). Three simulations of duration 3 μs were run for all the SDC helices. During each simulation, the two helices diffused randomly in the bilayer, encountered one another, and formed a dimer (Fig S1 right).

In our simulations, all four syndecan TMDs can form stable homo-dimers in which the helices were packed in a right-handed fashion with a negative value of the helix crossing angle of ~25° and interhelix separation of ~6 Å (centroid distance). The binding energies and the RMSD analysis showed that among the four homodimers, the SDC2 dimer is the most stable while SDC1 dimers are the least stable. In addition to forming the stable dimers with the helix crossing angle of ~25°, SDC1 helices can also form unstable dimers during the simulation processes with crossing angle of ~0°, i.e. parallel dimers, which have longer centroid distances (~9 Å) and smaller binding energies (~-308 kJ/mol), compared with the relatively stable right-handed model of SDC1 dimer (centroid distance of ~6 Å and binding energy of ~-311kJ/mol). This unstable model indicates the possibility of a competing conformation during the dimerization process.

To further analyze the molecular mechanisms of the dimerization, we examine the residue contact maps shown in Fig 2 and Fig S2 and the dimerization interfaces of the four dimers in Table 1. The SDC2 forms the most stable dimer with the “X” shape with the conserved dimerizing motif G8XXXG12 on one helix at the contact interface interacting with A7XXXI11 motif on the other helix (Fig 2C). Almost all the stable dimers share a similar conformation. This asymmetric arrangement of two helices was also found in the structures of the homodimer of SDC4-TM dimer in solution NMR studies[30]. The GXXXG motif is considered a common motif in mediating the homo-dimerization[31]. We propose that the G8XXXG12 located in the middle of the helix facilitates the formation of the “X”-shaped dimer, which maximizes the number of contacts on the interface with G8XXXG12 at the crossing and stabilizes the dimerization.

**Table 1.**
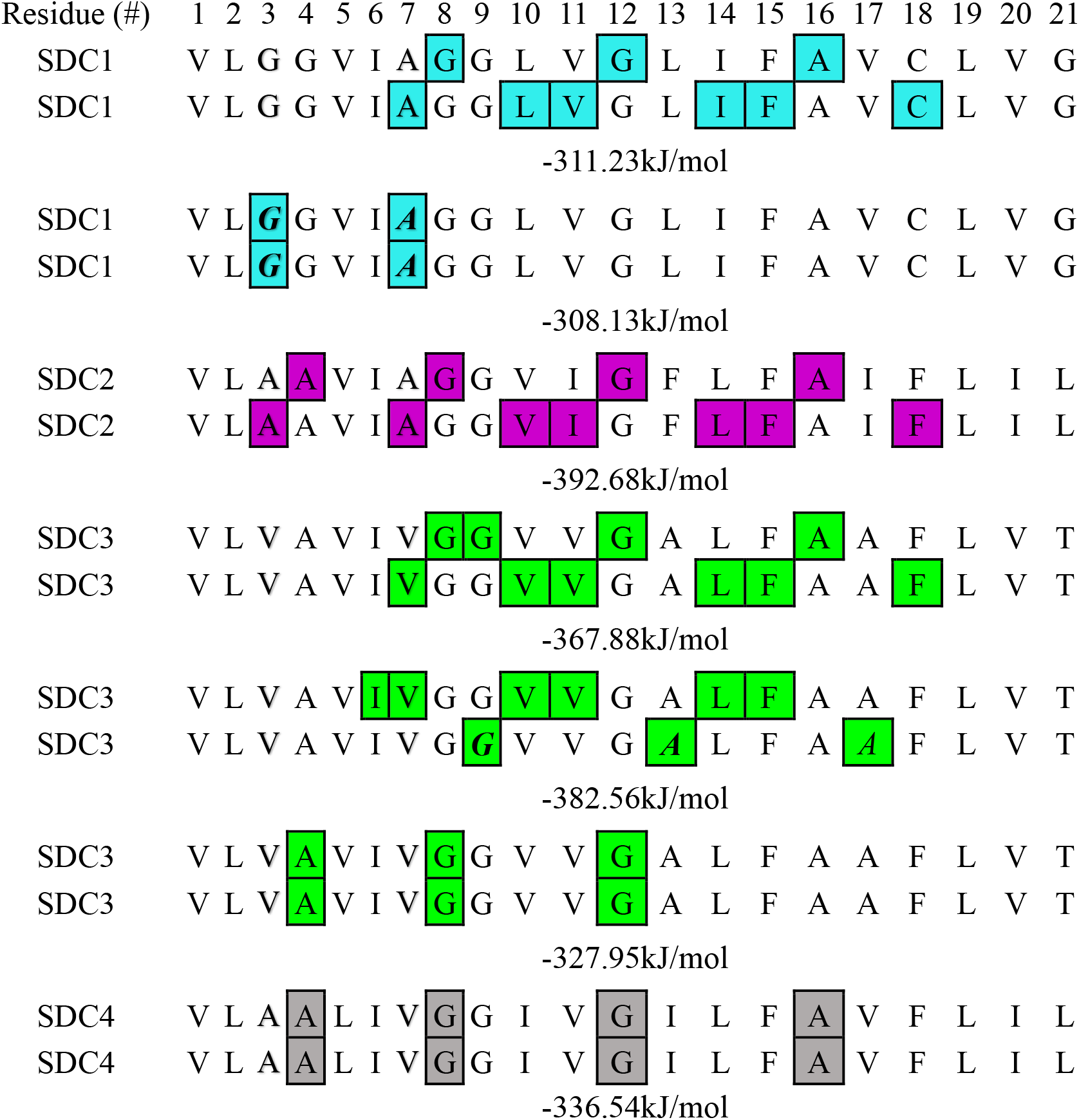
The contact residues and binding energy of sydecans homodimer

In addition to GXXXG motif, GXXXA motif, another common dimerizing motif in TMD dimerization, also appear in the contact interface in the dimers of SDC1 and SDC3. In SDC3 dimers (Fig S2C), the G9XXXA13 in the middle of the helix facilitates the formation of “X”-shaped dimer and it is even more stable (Fig 1C top, −382.56kJ/mol) than the dimer with G8XXXG12 motif on the contact interface (Fig 1C top, −367.88kJ/mol). As we mentioned above, SDC1 can form an unstable parallel dimer, in which, the residues Gly3, Ala7 of both TMD helices formed the helix/helix interface G3XXXA7 (Fig 2A c) while the conserved G8XXXG12 motif forms the interface in the right-handed model of SDC1 dimer (Fig 2B c). The G3XXXA7 motif in SDC1 locates far away from the middle of the helix, which facilitates the formation of the “V”-shaped dimer with the G3XXXA7 on the contact interface. This results in a significantly reduced number of contacts on the interface when compared with “X”-shaped dimer, making the “V”-shaped dimer unstable. Nonetheless, the existence of the “Y”-shape dimer competes with the formation of “X”-shaped dimer, resulting in the least stability of the dimerization of SDC1 among all the homodimers of SDCs. Our simulations are in good agreement with the experimental results reported by Dews et al. that each syndecan member has variable affinities for dimerization or oligomerization: SDC2 dimerizes more actively as compared to those of SDC3 and 4, while SDC1 dimerizes very weakly[31].

**Fig 1.**
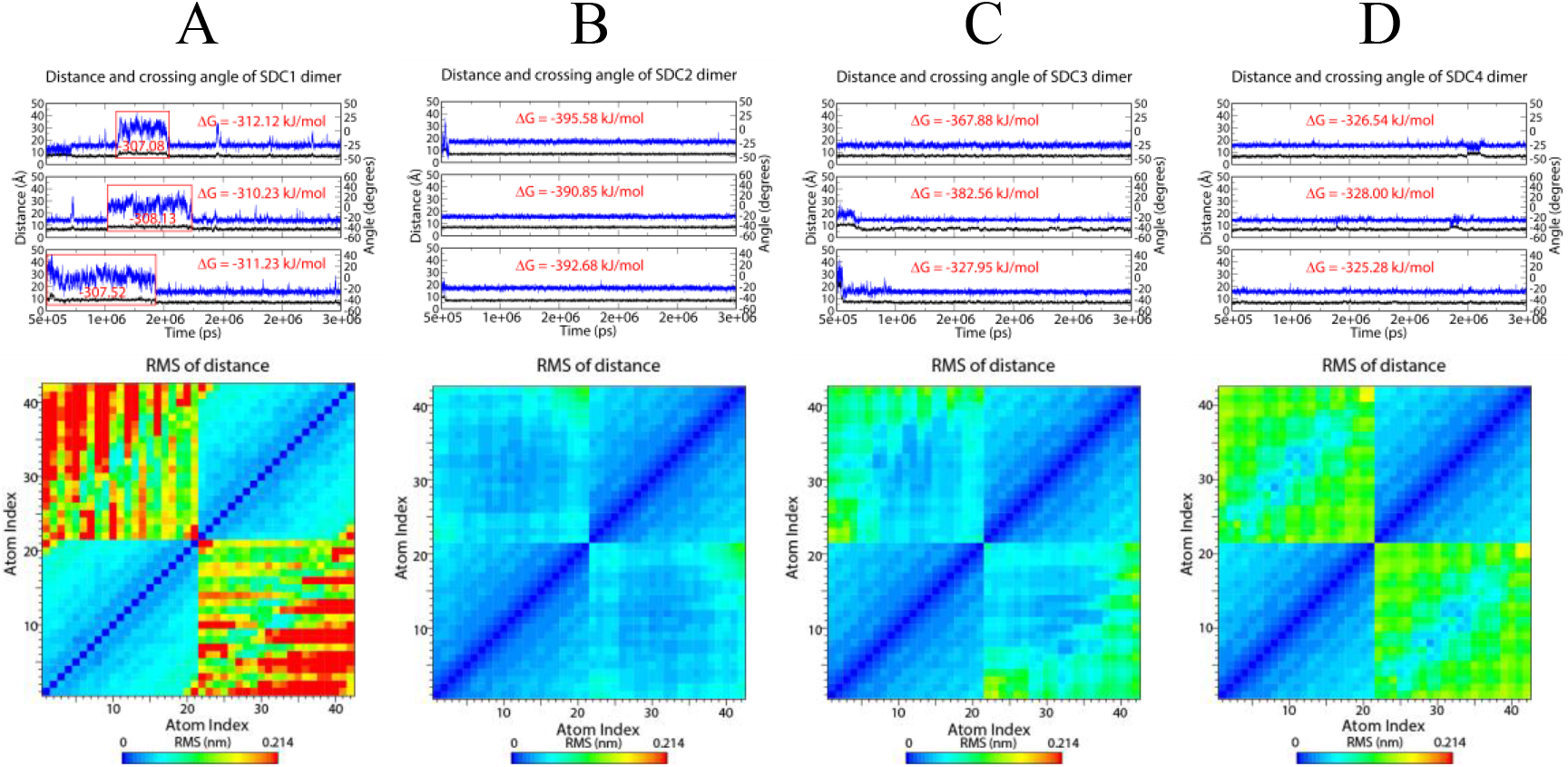
The stability of the dimers of syndecans TMDs. Top: The distance and crossing angle of two TMDs in the dimer during the simulation: the intersection angle (blue line), the distance (black line), and the g-mmpbsa binding energy are shown in red text. The unstable dimer of SDC1 was boxed in red with the corresponding binding energies. The total number of samples is 3. Bottom: RMSD of distance between beads on the dimer backbone.

**Fig 2.**
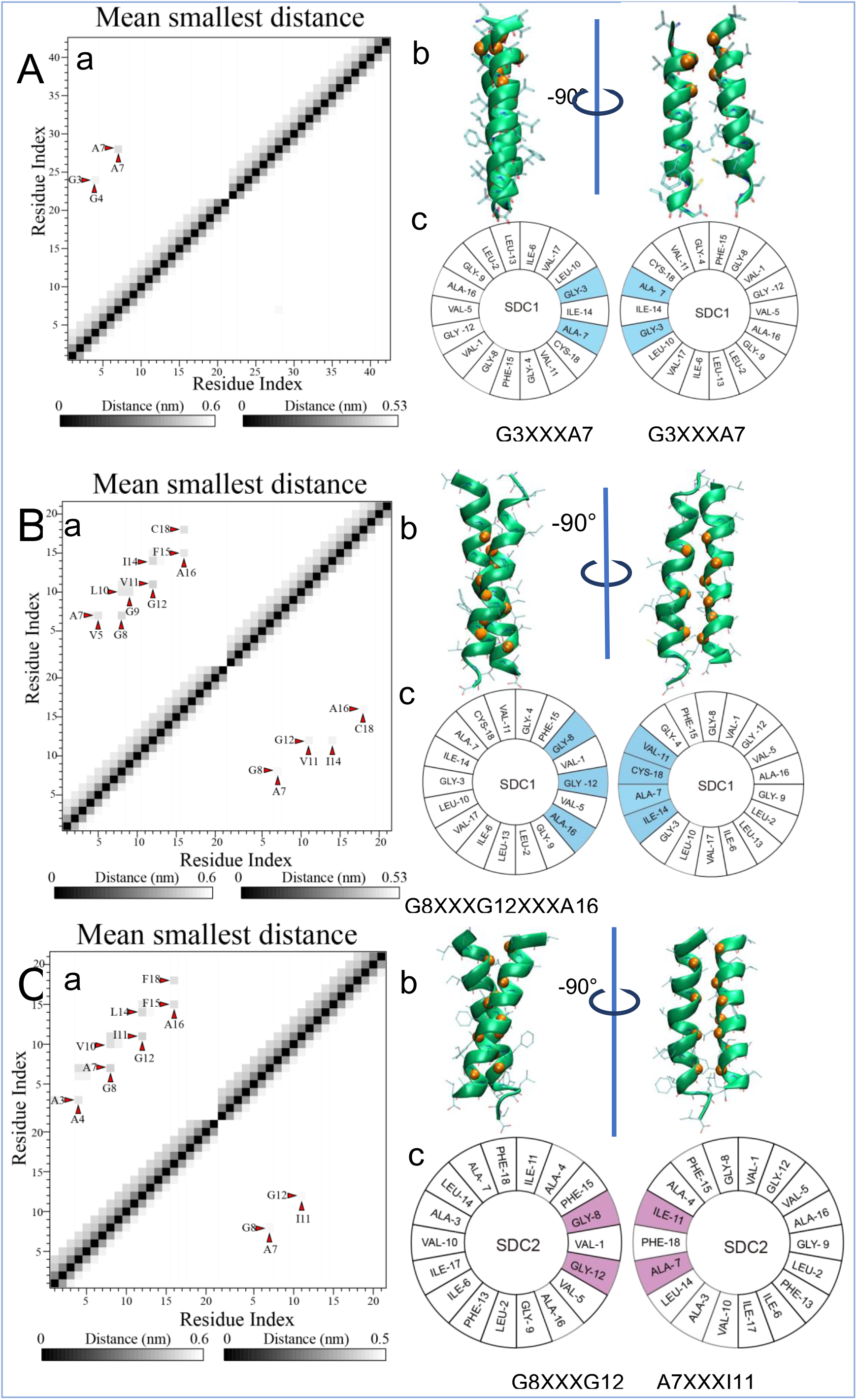
Contact interface of SDC1, SDC2 dimer. A. a, mean smallest distance of SDC1_1 dimer’s backbone bead (The small red triangle points to the darkest point, because the darker the color, the closer the distance) b. Cartoon pictures of SDC1 dimer. c. Helix wheel pictures B. SDC1_2. C. SDC2.

To further study how the GXXXA motif affects the stability of dimerization, a series of mutants were designed and the binding energies were calculated using G_mmpbsa. Wild type and mutant sequences are summarized in Table 2. To explore the aggregation stability of syndecans, the initial stable dimer structure experienced MD was used for PMF[32] calculations. The results from the PMF calculations are compared in Fig 3B in the form of free-energy profiles as a function of helix/helix distance. All four prologs of syndecan and their mutations are seen to have energy minima for associated, dimerized conformations, albeit with different well depths. SDC2 has the most depth free-energy minimum. SDC1 has the most shallow free-energy minimum. Our PMF results are similar to the results of g_mmpbsa binding free energy.

**Fig 3.**
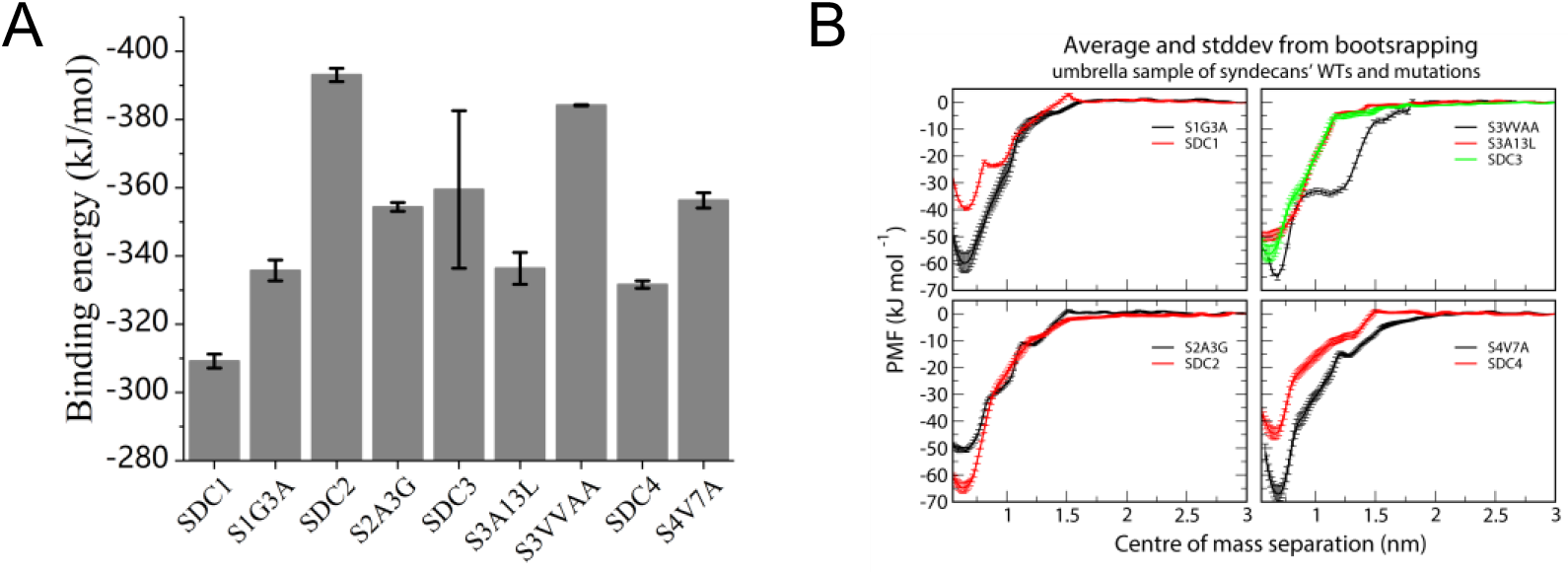
A. G_mmpbsa binding free energy of SDCs dimer WTs and mutations. B. Potential of mean force measured in umbrella sample simulations of SDCs dimer WTs and mutations. (red line represent the wild type, black line represent mutations)

**Table 2.** Syndecan WT TMD sequence and mutation TMD sequence

| Index | 1 | 2 | 3 | 4 | 5 | 6 | 7 | 8 | 9 | 10 | 11 | 12 | 13 | 14 | 15 | 16 | 17 | 18 | 19 | 20 | 21 |
| --- | --- | --- | --- | --- | --- | --- | --- | --- | --- | --- | --- | --- | --- | --- | --- | --- | --- | --- | --- | --- | --- |
| SDC1 | V<sup>255</sup> | L | <u>G</u> | G | V | I | <u>A</u> | G | G | L | V | G | L | I | F | A | V | C | L | V | G<sup>275</sup> |
| S1G3A | V | L | <u>A</u> | G | V | I | <u>A</u> | G | G | L | V | G | L | I | F | A | V | C | L | V | G |
| SDC2 | V<sup>155</sup> | L | <u>A</u> | A | V | I | <u>A</u> | G | G | V | I | G | F | L | F | A | I | F | L | I | L<sup>175</sup> |
| S2A3G | V | L | <u>G</u> | A | V | I | <u>A</u> | G | G | V | I | G | F | L | F | A | I | F | L | I | L |
| SDC3 | V<sup>385</sup> | L | <u>V</u> | A | V | I | <u>V</u> | G | <u>G</u> | V | V | G | <u>A</u> | L | F | A | A | F | L | V | T<sup>405</sup> |
| S3A13L | V | L | V | A | V | I | V | G | <u>G</u> | V | V | G | <u>L</u> | L | F | A | A | F | L | V | T |
| S3VV-AA | V | L | <u>A</u> | A | V | I | <u>A</u> | G | G | V | V | G | A | L | F | A | A | F | L | V | T |
| SDC4 | V<sup>150</sup> | L | <u>A</u> | A | L | I | <u>V</u> | G | G | I | V | G | I | L | F | A | V | F | L | I | L<sup>170</sup> |
| S4V7A | V | L | <u>A</u> | A | L | I | <u>A</u> | G | G | I | V | G | I | L | F | A | V | F | L | I | L |

\*Underlined marks indicate residues before and after the mutation.

We designed two mutants S1-G3A and S3-A13L to remove the G3XXXA7 in SDC1 and G9XXXA13 in SDC3, respectively. It was found that for both S1-G3A and S3-A13L form stable dimers with a single interface at the conserved G8XXXG12 motif without the competing motif GXXXA. In addition, The S2-A3G mutant with G3XXXA7 showed lower stability than the WT-SDC2 with A3XXXA7 (Fig S4B). Thus, the competing effect of G3XXXA7 motif disturbs the dimerization of SDC TMDs through the interface at G8XXXG12 motif. Furthermore, the S3-VVAA and S4-V7A mutants showed higher stabilities than the WT SDC3 and WT SDC4, respectively, indicating that the A3XXXA7 motif may stabilize the dimerization. These results agree with our hypothesis that dimerizing motif G3XXXA7 far away from the middle of the helix facilitates an unstable competing conformation and decreases the stability of dimerization.

### The heterodimerization of SDC TMDs

In addition to homodimerization, heterodimerization of SDC TMDs was performed using a similar method of MD simulations and data analysis including listing the residues involved in the dimeric interface and their g_mmpbsa binding energy in the table (Table 1). Putting together the free energy of the homodimer and the heterodimer, a three-dimensional map (Fig 4A) was made. Based on the magnitude of these energies, a table was made to show the relative strength of the dimer (Fig 4B). Our results showed that SDC1 and SDC2 have the weakest and strongest tendency to form heterodimers with other SDCs, respectively. The most stable heterodimers are SDC2-3 and SDC2-4. However, multiple conformations with large energy differences were found in most heterodimers, indicating the complexity of heterodimerization of SDCs.

**Fig 4.**
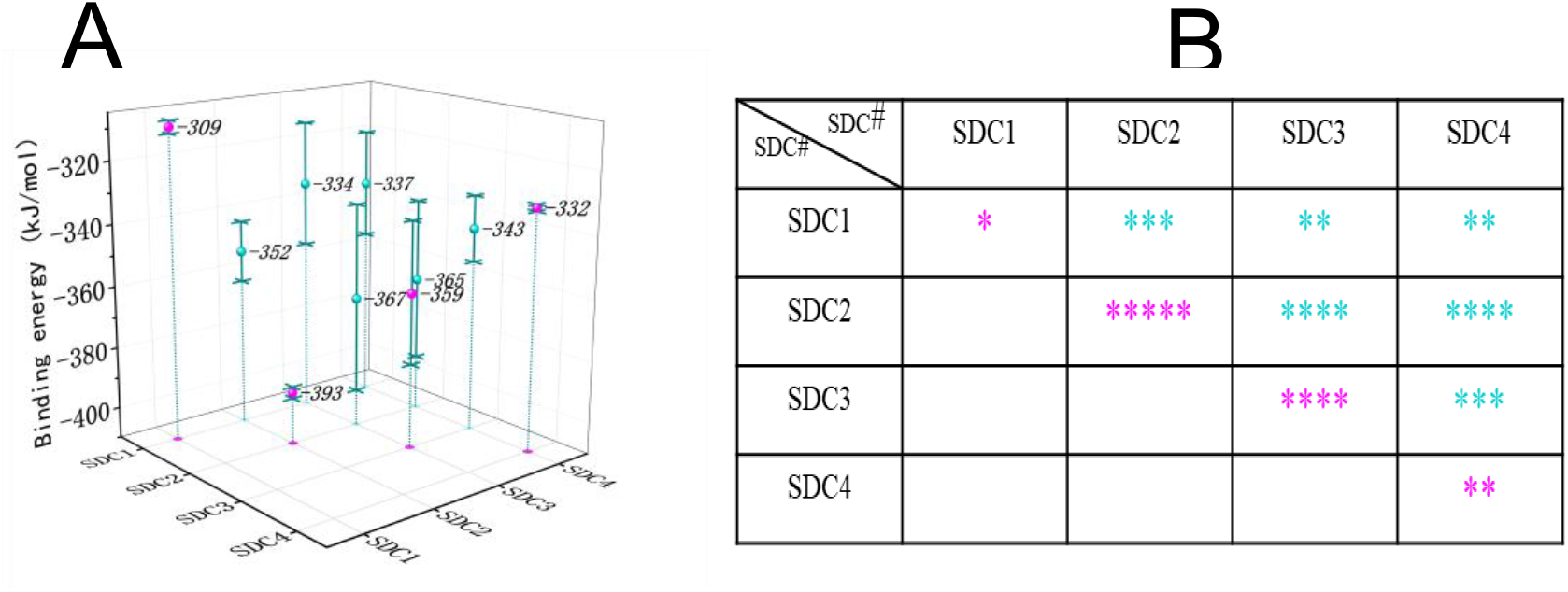
Relative strengths of syndecan TMD interactions. A. Binding free energy of syndecans dimer(including both homodimer and heterodimer) B. a table listing relative strengths of syndecan TMD dimer stability. The more asterisks, the stronger the interaction. Magenta—homodimer, cyan—heterodimer.

To clearly demonstrate the dimeric interface of the heterodimer, we here primarily show two conformations in the lowest energy SDC23 and one conformation in SDC24. As shown in Fig 5, both of G8XXXG12 and G9XXXA13 motifs from SDC3 can form the contact interface in the dimers of SDC2-3 while G8XXXG12 from SDC4 is presented in the contact interface from the dimer. However, the A7XXXI11 Instead of G8XXXG12 from SDC2 is presented in the dimeric interface in the heterodimers SDC2-3 and SDC2-4. In addition, we suspect that the multiple contact interfaces of SDC2 and SDC3 in the heterodimers can be used for further oligomerization of SDC2 and SDC3. Our simulations showed that the heterodimer of SDC2-3 can form a heterotrimer with either one SDC2 helix or one SDC3 helix (Figure S4). In fact, the experimental study [31] has shown that SDC2 can form a trimer with two SDC3s.

**Fig 5.**
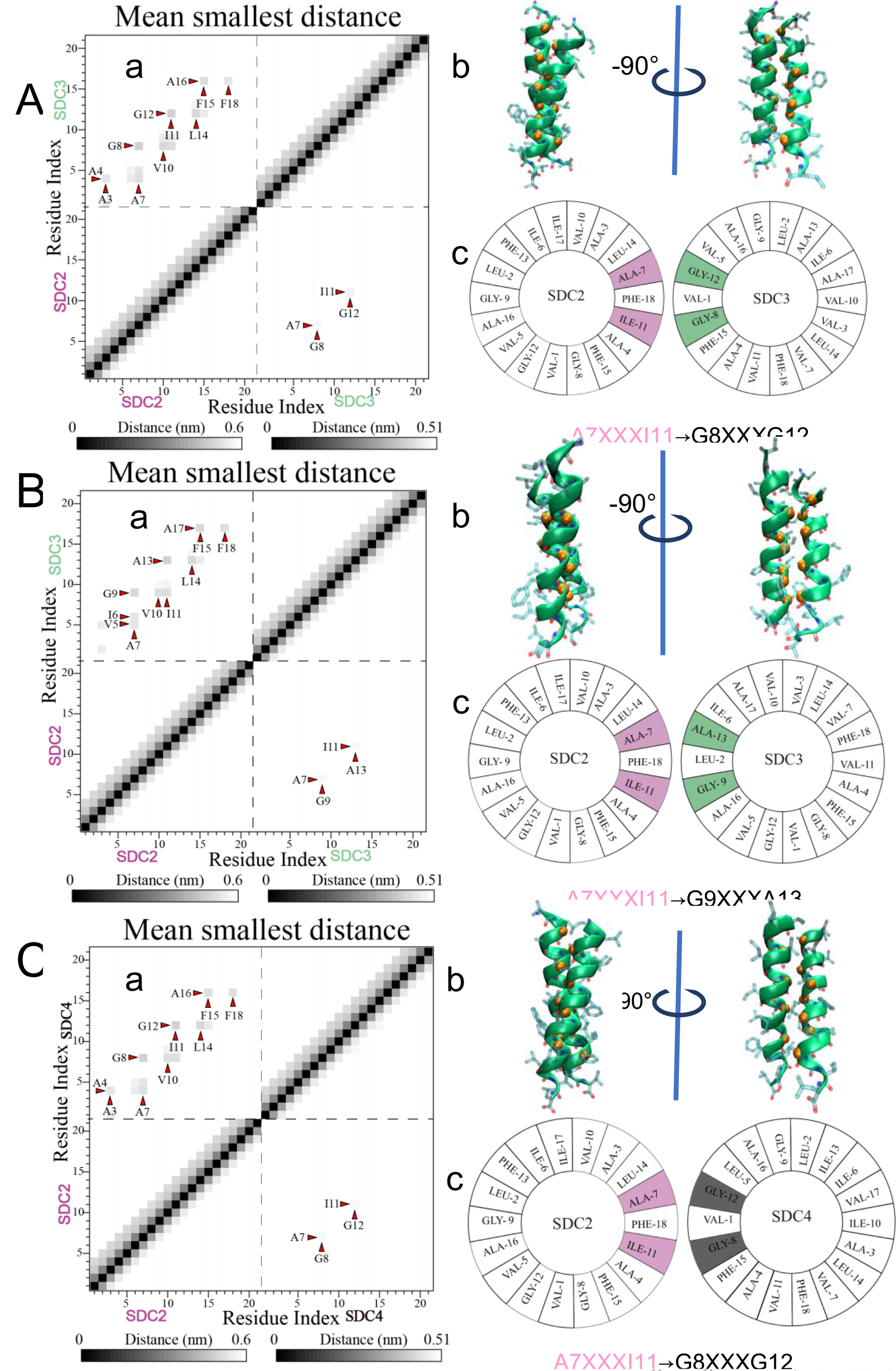
A Contact interface of SDC23, SDC24 dimer. A. a, mean smallest distance of SDC23_1 dimer’s backbone bead(The small red triangle points to the darkest point, because the darker the color, the closer the distance) b. Cartoon pictures of SDC23_1 dimer. c. Helix wheel pictures B. SDC23_2. C. SDC24.

## Conclusions

Both GXXXG and GXXXA are considered common motifs in mediating the dimerization of TMDs. In this study, we found that the GXXXG and GXXXA motifs in the middle of the SDC TMD helix facilitated the formation of an X-shaped dimer, which maximized the number of contacts at the interface and thus stabilized the dimerization. However, because G3XXXA7 in SDC1 is far away from the middle of the helix, it facilitated the formation of a Y-shaped dimer, which results in a reduced number of contacts at the interface and thus destabilize the dimerization. Multiple dimerizing motifs and contact interfaces in the dimers of SDCs results in the complexity of the homo- and hetero-dimerization. Our results shed lights on the molecular mechanism of the dimerization of TMDs and the importance of the locations of the multiple dimerizing motifs.

## Author Contributions

Jialin Chen designed this study, performed the simulations, analyzed the data, and wrote the article. Fengli Wang performed the simulations and analyzed the data. Chengzhi He and Shizhong Luo designed this study, analyzed the data and revised the article.

## Acknowledgment

This work was supported by the National Natural Science Foundation of China (21672019, 21372026, 21402006) and the Fundamental Research Funds for the Central Universities (Grand No. XK1701). This work was partly supported by CHEMCLOUDCOMPUTING.

